# Genome-wide association study of host resistance to bacterial leaf streak in a subset of the world barley core collection

**DOI:** 10.1101/2025.10.31.685662

**Authors:** Diel Donne Velasco, Gongjun Shi, Robert Bruggeman, Richard Horsley, Thomas Baldwin, Zhaohui Liu

## Abstract

The bacterial leaf streak (BLS) disease of barley, caused by *Xanthomonas translucens* pv. *translucens* (Xtt), has become increasingly important worldwide in recent years. Inefficacy of chemical control methods leaves deployment of host resistance to be the only option to manage this disease. However, current commercial varieties are mainly susceptible to BLS. Therefore, our goal was to identify sources of resistance from diverse barley germplasms and map associated genetic factors. To do so, we evaluated a subset of the World Barley Core Collection (BCC), consisting of 198 accessions, on their reaction to BLS from 2013 to 2016 under natural or artificially inoculated disease pressures. Ten accessions exhibited consistently low disease severities over four years of evaluations. Using genotype data from the T3/Barley database, genome-wide association studies were conducted to identify marker-trait associations (MTAs) in this barley mini-core panel for BLS resistance. Utilizing four mixed-model analyses (MLM, MLMM, FarmCPU, BLINK), five significant MTAs were consistently identified from at least two mixed model analyses including two in chromosome 2H, and one each in chromosomes 5H, and 7H. Associations in chromosomes 2H and 5H appear to be in the same region with loci identified in a previous association study, reinforcing their potential relevance. The identified resistant barley accessions and associated markers will be valuable inbreeding BLS-resistant barley varieties.

**Core ideas:**

– A subset of world barley collection was used in the identification of sources of resistance and marker-trait association for resistance for bacterial leaf streak.
– Ten accessions consistently showed low disease severity across different years, highlighting their value in BLS resistance breeding.
– Five MTAs were consistently identified using different models, with two aligning with previously identified resistance QTLs.

## 1 INTRODUCTION

Bacterial leaf streak disease (BLS) of cereals, caused by the several pathovars of *Xanthomonas translucens* (Xt), has become increasingly important in the past years for both wheat and barley. Although the disease was first reported on barley in 1917 by Jones et al., efforts to understand this pathosystem have progressed slowly, potentially due to the sporadic and low incidence of the disease. However, BLS disease in both barley and wheat has been steadily increase in the upper Midwestern United states, specifically in the fields of North Dakota, Minnesota, and South Dakota, excluding the drought years of 2017 and 2021 (Adhikari et al. 2011; Kandel et al. 2012; Curland et al. 2018; Friskop et al. 2023). The persistence of BLS in barley and wheat fields across major cereal production states has elevated the importance of the disease.

As BLS affects the leaves that sustain crop productivity, yield parameters including test weight, percent small seed, and total yield have been observed to be directly affected by BLS infection in both barley and wheat (Shane et al. 1987; Tillman et al. 1996). Shane et al. (1987) reported that 50% disease severity on the flag leaf resulted in an 8-13% reduction in kernel weight for both cereal crops, while a 100% disease severity on the flag leaf incurred 13-34% loss on kernel weight. These reductions were evident with the loss of grain plumpness on severely infected plants. In addition, they also reported that the grain protein content in wheat was positively correlated with disease severity, suggesting a potential impact on dough elasticity. In wheat, compatible Xt pathovars cause two diseases - bacterial leaf streak and black chaff diseases. Although both may impact the reproductive stage of wheat, Tillman et al. (1994) indicated that only bacterial leaf streak severity relates to yield loss but not black chaff severity. Moreover, they found that bacterial leaf streak and black chaff severities in winter wheat cultivars does not correlate consistently, indicating that one disease phenotype cannot be used to screen for the other (Tillman et al. 1996).

Recent estimations on 44 hard red spring wheat (HRSW) varieties have shown yield reductions of up to 60% (Friskop et al. 2023). Although current yield loss estimates for infected barley have not been determined, the potential 34% reduction in kernel weight reported by Shane et al. (1987), provides a strong indication of the negative impact on productivity, grain quality, and seed viability. This highlights the need to identify effective management strategies for controlling the bacterial leaf streak disease in barley.

The limited understanding of the pathogen’s biology, epidemiology, and mechanism of host interactions poses a significant challenge for researchers in developing effective BLS management strategies. With the pathogen being seed-borne (Duvelier et al. 1997; Jones et al. 1917), seeds were considered as primary sources of inoculum, making seed treatment a practical option for disease control. However, assessment of seed treatments and foliar pesticide application had shown significant impact on seed viability and inconsistency in efficacy to lower disease levels in the field (Forster and Schaad 1988; Sands et al. 1989; Duveiller et al. 1997; Lux et al. 2023).

Several studies in barley and wheat have been conducted to characterize varietal reactions to BLS with the aim of identifying donors of resistance and mapping the corresponding resistance genes (Duveiller et al. 1993; Tillman et al 1996; El Atarri et al. 1998; Tillman and Harrison 1996; Adhikari et al. 2012; Gurung et al. 2014; Ritzinger et al 2023a, b). In barley, El Atarri et al. (1998) evaluated the genetic variability of doubled-haploid lines for BLS resistance, identifying two QTLs on chromosome 3H and one on chromosome 7H, with multilocus allelic effects explaining approximately 30% of the observed phenotypic variation. In a more recent study to identify potential sources of resistance to BLS, Ritzinger et al. (2023a) conducted field evaluations in two years on a total of 2,094 accessions that belong to barley improvement programs from the University of Minnesota (UMN) and Anheuser-Busch InBev (ABI), landrace panels from Leibniz Institute of Plant Genetics and Crop Plant Research (IPK) and Ethiopian Eritrean Barley Collection (EEBC), and introgression lines from wild barley in the genetic background of cultivar ‘Rasmusson’. Thirty-two accessions showed consistent BLS resistance that are within a similar maturity range of standard barley cultivars commonly planted in the Upper Midwest United States. In a following work, Ritzinger et al. (2023b) conducted a GWAS on two barley breeding panels to uncover BLS resistance markers. Their study identified one significant association in the first panel and seven significant associations in the second, with resistance-linked markers located on chromosomes 1H, 2H, 3H, 5H, 6H and 7H. Interestingly, one of the associated markers identified was linked to the photoperiod response gene *Ppd-H1* on chromosome 2H. This finding supports the hypothesis that plant maturity influences the observed BLS response in barley.

Genotyping platforms and services have made it possible to conduct GWAS using thousands to millions of SNPs, facilitating the discovery of QTLs of interest and identifying resistance sources across diverse populations. Higher genetic diversity within the population increases the likelihood of capturing a wider range of genetic variants and enhances the chances of detecting significant associations between markers and traits of interest. The Barley Core Collection (BCC) from USDA-National Small Grains Collection that represents the global diversity of barley was established to identify alleles that will be valuable in developing adaptable varieties to climate-related stresses. The BCC consists of 2,417 accessions that have been genetically characterized and have been evaluated for several agronomic and disease resistance traits (Muñoz-Amatriain et al. 2014). Recently, a subset of this collection, designated barley mini-core panel, has been successfully used to identify MTAs for barley resistance to hessian fly (Karki et al. 2024). In a similar approach, we conducted GWAS to identify genomic regions that are associated with BLS resistance using this mini-core set of the BCC.

## 2 MATERIALS AND METHODS

### 2.1 Mini-core panel accessions

The mini-core panel evaluated in this study is a subset of the USDA-NSGC Core Collection. This panel consists of 198 accessions, including 70 landraces, 59 cultivars, 36 breeding lines, and 32 accessions with unknown improvement status. These accessions represent a broad geographical diversity, sourced from 69 countries worldwide (Muñoz-Amatriaín et al. 2014). All accessions are a spring type, with 88 being two-rowed types and 110 six-rowed types. Further details on each accession are provided in Supplementary Table 1.

### 2.2 Phenotypic data and analysis

The mini-core panel was evaluated on field either under natural disease pressure and/or artificially inoculated nursery from 2013 to 2016 in Fargo, ND. Evaluations in 2013 and 2015 were taken from natural disease pressure while evaluation in 2016 was done from an artificially inoculated nursery. Evaluations in 2014 were carried out in both conditions. The evaluations were designated as 2013NAT, 2014NAT, 2014ART, 2015NAT and 2016ART. For all evaluations, ‘Pinnacle’ was used as a susceptible check. For evaluations under natural disease condition, the genotypes were planted as a single row in the field with a length of four feet without replication. For artificially inoculated disease conditions, the genotypes were planted in hill plot (1 x 1 ft2 for each entry) with four replicates following randomized complete block design. Bacterial inoculums were prepared from a local Xtt strain B2 (Peng et al. 2016). To prepare inoculum, the bacterial strain was cultured on WBA media (Wen et al. 2017) for two days and used to make a suspension at a concentration of 1 x 10^8^ colony forming units. The bacterial suspension was subsequently mixed with carborundum and was sprayed onto test entries using a gas-powered backpack sprayer. The inoculations were performed at the late tillering stage following the procedure described in Green et al. (2023). For all four years, disease scorings were performed at the soft dough stage based on a 1-9 scale that was used for wheat BLS (Green et al. 2023).

For GWAS, Given the unbalanced data across five environments the best linear unbiased predictors (BLUP) for the disease response were calculated in R using the packages lme4 (Bates et al., 2015) and lmerTest (Kuznetsova et al., 2017). BLUPs were estimated using a mixed-effect model, where year, taxa, their interaction (year:taxa), and replicates nested in years (year/rep) were treated as random effects. The estimated BLUP values were subsequently used for association mapping.

### 2.3 Genotypic data and analyses

The BCC accessions were genotyped using the 9k Illumina Infinium iSelect SNP assay (Muñoz-Amatriain et al. 2014). The genotypic data were retrieved from the Triticeae Toolbox/Barley (https://barley.triticeaetoolbox.org/). Markers with a minor allele frequency (MAF) of less than 0.05 were excluded using TASSEL v5 (Bradbury et al., 2007). A total of 5,568 SNP markers were obtained for the association mapping analysis.

### 2.4 Population structure and linkage disequilibrium analyses

An inference on the population structure of the mini-core panel was carried out using STRUCTURE v2.3.4 with the admixture model (Pritchard et l., 2000). The burn-in period was set at 5,000 with simulations being conducted at 50,000 replications and iterated 10 times. The range of assumed sub-populations (K) was set from 1 to 10. Simulation results were collated using STRUCTURE HARVESTER (Earl and vonHoldt, 2005), which was used to subsequently identify the best-fitted number of K within the mini-core panel. Admixture in the population was accounted for based on the ancestry plot that employed 70% membership coefficient.

Linkage disequilibrium estimates, r2, were generated using TASSEL v5 with a sliding window size of 50. All LD estimates for each chromosome were plotted against their respective physical distances and were visualized in RStudio (RStudio Team, 2020). Similarly, mean r^2^ values across all chromosomes were calculated and visualized against physical distances in RStudio.

### 2.5 Association mapping and multiple-comparison correction

To investigate potential significant marker-trait associations (MTAs), GWAS was performed in R using the package GAPIT 3.4 (Wang and Zhang, 2021). Multiple models were used for the analysis, including Mixed Linear Model (MLM), Multi-Locus Mixed Model (MLMM), Fixed and random model Circulating Probability Unification (FarmCPU), and Bayesian-information and Linkage-disequilibrium Iteratively Nested Keyway (BLINK), to identify consistent marker-trait associations.

In the GWAS models, the population structure (Q) and kinship (K) were used as covariates to account for confounding effects. The Q-matrix was derived from the optimal number of subpopulations (k) identified within the population, while the K-matrix was calculated using the VanRaden algorithm through GAPIT 3.4 (VanRaden, 2008). The Li and Ji method (2005) for multiple testing correction was used to determine the significance threshold.

## 3 RESULTS

### 3.1 Phenotypic Evaluation

Across all the evaluations, the susceptible check ‘Pinnacle’ showed high levels of susceptibility (disease severity score =8.0). The reaction of accessions in the mini-core panel ranged from highly resistant (severity score =1) to highly susceptible (severity score=9.0) and the distribution of accessions, in each category, for each environment were shown in Figure 1A. The average disease means were 3.6, 3.8, 3.6, 3.2 and 4.2 for 2013NAT, 2014NAT, 2014ART, 2015NAT and 2016ART, respectively. Based on the computed BLUP values across five environments, the population is composed of 25 resistant (< 3.5), 109 intermediate (3.5 to 5.5), and 64 susceptible (≥ 5.5) accessions (Figure 1B). Twenty-one genotypes showed consistent low severity scores across all environments (Table 1).

**Figure 1.**
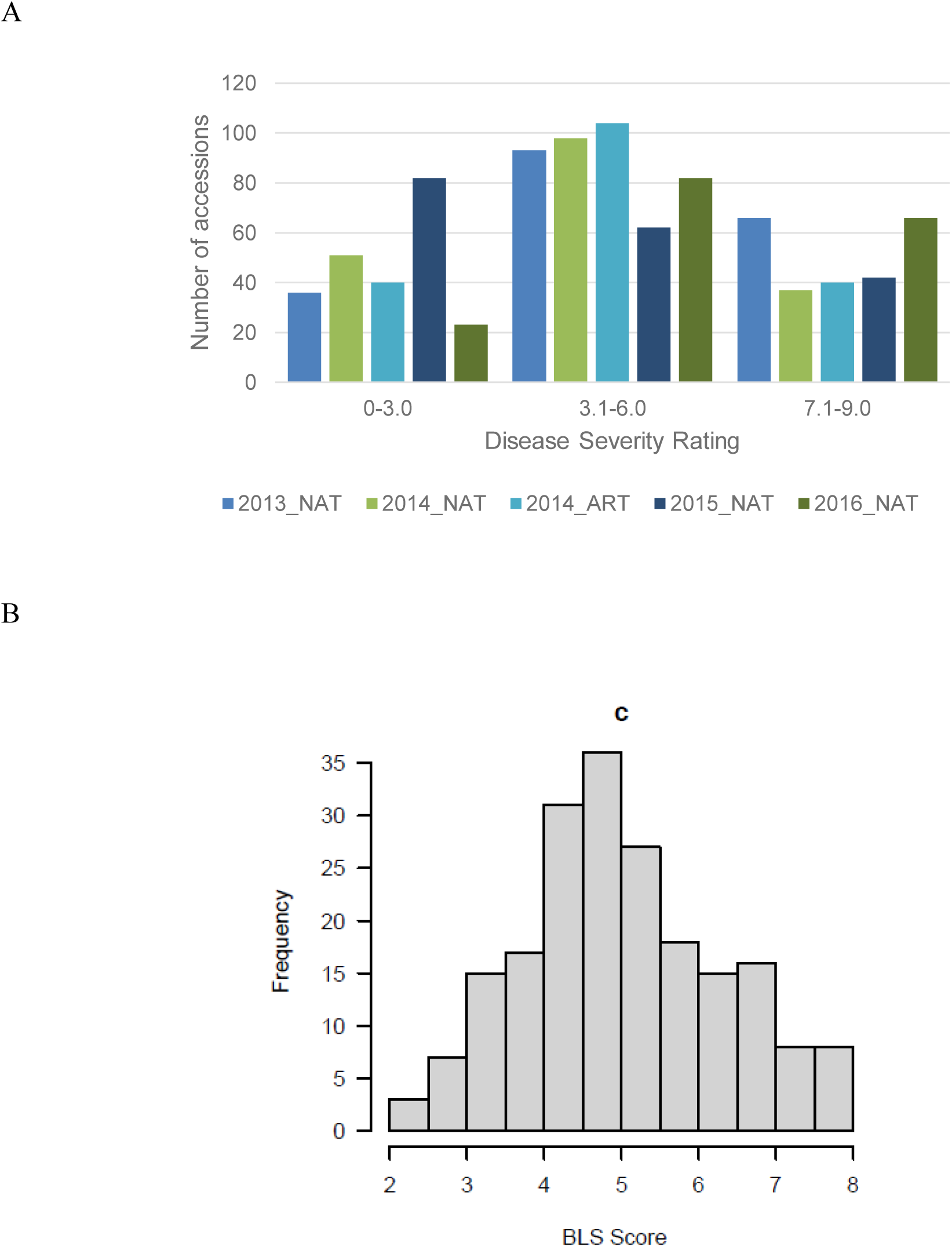
(A) Distribution of BLS severity rating across years and environments (NAT – natural infection; ART – artificially inoculated); (B) Distribution of calculated BLS severity ratings (BLUP) on the mini-core panel from the data collected in 2013 to 2016.

**Table 1.**
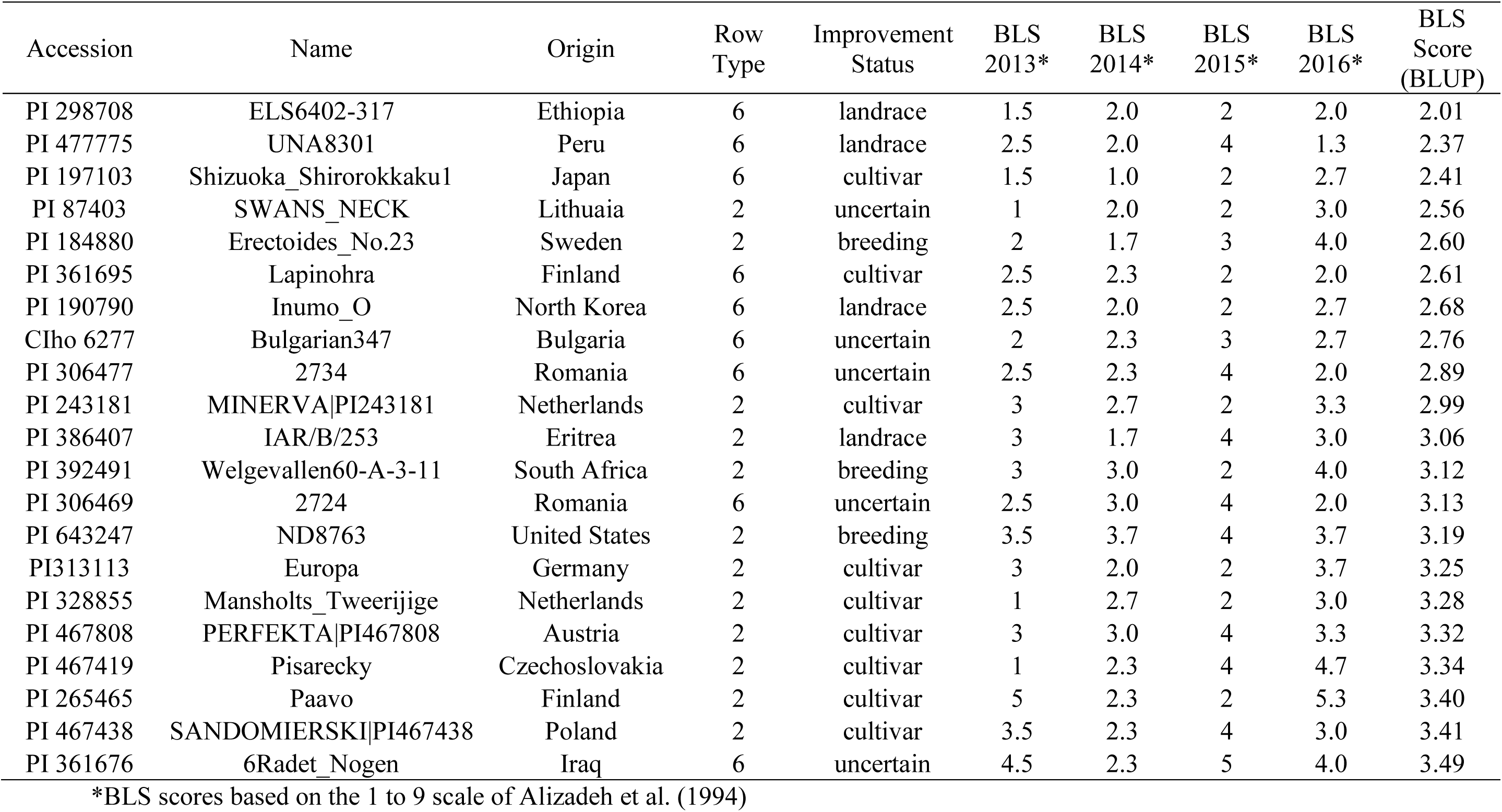
Accessions that have consistent low infections during the trials conducted in 2013 to 2016.

| Accession | Name | Origin | Row Type | Improvement Status | BLS 2013* | BLS 2014* | BLS 2015* | BLS 2016* | BLS Score (BLUP) |
| --- | --- | --- | --- | --- | --- | --- | --- | --- | --- |
| PI 298708 | ELS6402-317 | Ethiopia | 6 | landrace | 1.5 | 2.0 | 2 | 2.0 | 2.01 |
| PI 477775 | UNA8301 | Peru | 6 | landrace | 2.5 | 2.0 | 4 | 1.3 | 2.37 |
| PI 197103 | Shizuoka_Shirorokkaku1 | Japan | 6 | cultivar | 1.5 | 1.0 | 2 | 2.7 | 2.41 |
| PI 87403 | SWANS_NECK | Lithuania | 2 | uncertain | 1 | 2.0 | 2 | 3.0 | 2.56 |
| PI 184880 | Erectoides_No.23 | Sweden | 2 | breeding | 2 | 1.7 | 3 | 4.0 | 2.60 |
| PI 361695 | Lapinohra | Finland | 6 | cultivar | 2.5 | 2.3 | 2 | 2.0 | 2.61 |
| PI 190790 | Inumo_O | North Korea | 6 | landrace | 2.5 | 2.0 | 2 | 2.7 | 2.68 |
| CIho 6277 | Bulgarian347 | Bulgaria | 6 | uncertain | 2 | 2.3 | 3 | 2.7 | 2.76 |
| PI 306477 | 2734 | Romania | 6 | uncertain | 2.5 | 2.3 | 4 | 2.0 | 2.89 |
| PI 243181 | MINERVA\|PI243181 | Netherlands | 2 | cultivar | 3 | 2.7 | 2 | 3.3 | 2.99 |
| PI 386407 | IAR/B/253 | Eritrea | 2 | landrace | 3 | 1.7 | 4 | 3.0 | 3.06 |
| PI 392491 | Welgevallen60-A-3-11 | South Africa | 2 | breeding | 3 | 3.0 | 2 | 4.0 | 3.12 |
| PI 306469 | 2724 | Romania | 6 | uncertain | 2.5 | 3.0 | 4 | 2.0 | 3.13 |
| PI 643247 | ND8763 | United States | 2 | breeding | 3.5 | 3.7 | 4 | 3.7 | 3.19 |
| PI313113 | Europa | Germany | 2 | cultivar | 3 | 2.0 | 2 | 3.7 | 3.25 |
| PI 328855 | Mansholts_Tweerijge | Netherlands | 2 | cultivar | 1 | 2.7 | 2 | 3.0 | 3.28 |
| PI 467808 | PERFEKTA\|PI467808 | Austria | 2 | cultivar | 3 | 3.0 | 4 | 3.3 | 3.32 |
| PI 467419 | Pisarecky | Czechoslovakia | 2 | cultivar | 1 | 2.3 | 4 | 4.7 | 3.34 |
| PI 265465 | Paavo | Finland | 2 | cultivar | 5 | 2.3 | 2 | 5.3 | 3.40 |
| PI 467438 | SANDOMIERSKI\|PI467438 | Poland | 2 | cultivar | 3.5 | 2.3 | 4 | 3.0 | 3.41 |
| PI 361676 | 6Radet_Nogen | Iraq | 6 | uncertain | 4.5 | 2.3 | 5 | 4.0 | 3.49 |

507 \*BLS scores based on the 1 to 9 scale of Alizadeh et al. (1994)

### 3.2 Population Structure and LD Decay

Population structure analysis showed peaks at Δk = 2 and Δk = 4, suggesting the presence of 2 or 4 subpopulations (Figure 2A). Consistent with previous population structure analysis conducted on the entire barley iCore collection, and other mini-core sets (Muñoz-Amatriaín et al. 2012; Karki et al. 2023), at Δk = 2, the population structure (Q) matrix effectively divided the population according to row type. To investigate the second peak at Δk = 4, a membership coefficient of >0.7 was used to assign accessions in the four subpopulations, while accessions below this threshold were considered admixed (Supplemental Figure S1B). Similar with the findings of Muñoz-Amatriaín et al. (2012), the accessions are grouped into their geographical origins (Figure 2). In the BLS mini-core panel, Subpopulation 1 consists of two-rowed cultivars and breeding lines primarily from Europe, with some accessions from New Zealand and South America. Subpopulation 2 is composed of mostly six-rowed landraces from Asia. Subpopulation 3 includes six-rowed accessions from Mediterranean, Central, and South American regions (Supplemental Table S1). Subpopulation 4 consists of six-rowed cultivars and breeding lines from Canada, the United States, and Europe, as well as landraces from Asia (Supplemental Table S1). About a third of the mini-core panel is admixed (Supplementary Table S2).

**Figure 2.**
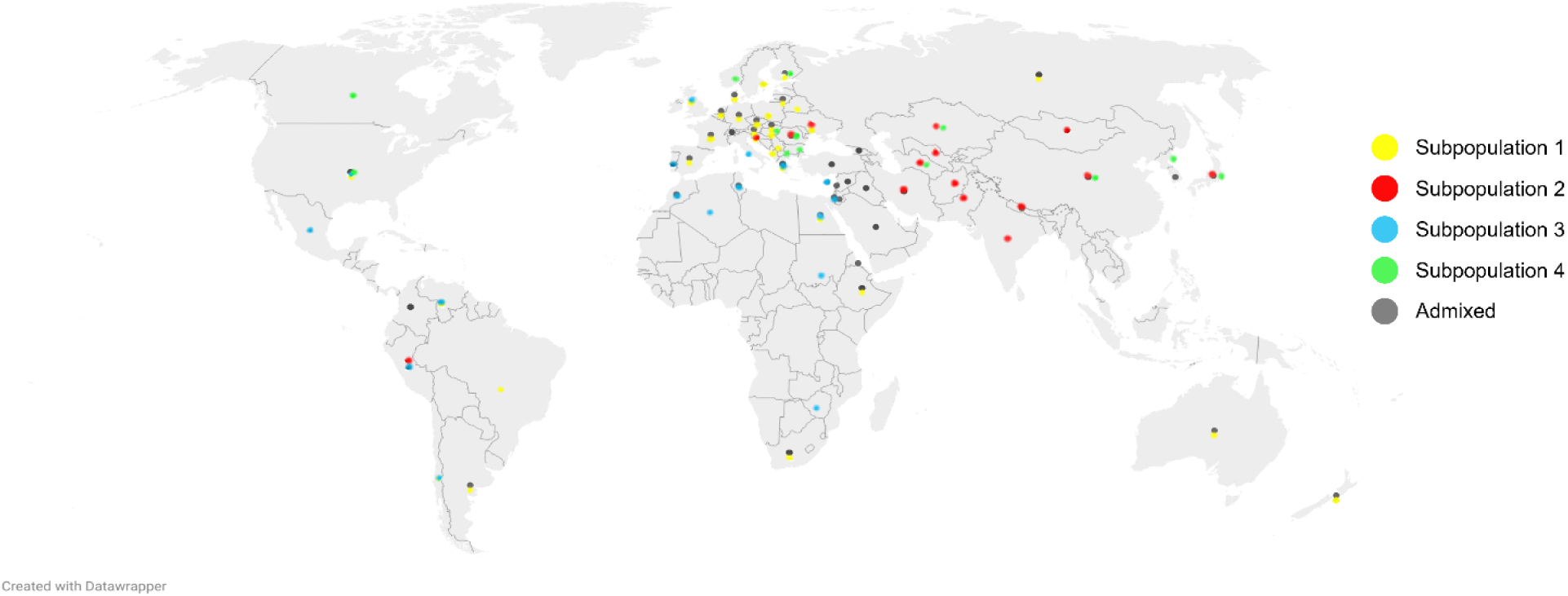
Geographical origins of accessions in each of the four subpopulations within the barley mini-core panel. Geographical origins of admixed accessions within the panel.

The overall LD of the mini-core panel exhibited a rapid decay pattern with increasing physical distance of markers. However, investigations on the LD decay for individual subpopulations revealed differences in their patterns. Based on threshold r^2^ = 0.1, rapid LD decay was observed in Subpopulations 2 (distance = 1.20 Mb) and 3 (distance = 0.49 Mb) while steadier decay patterns were observed in Subpopulations 1 (distance = 4.4 Mb) and 4 (distance = 4.4 Mb) (Supplemental Figure S2). The differences in LD decay patterns observed could likely reflect the composition of each subpopulation. Subpopulations 1 and 4 are predominantly composed of cultivars and breeding lines that imply that these accessions have been subjected to constant positive selection for favorable traits. This could potentially have led to lower genetic variability, within the subpopulation, and could explain the steady LD decay patterns each exhibited. In contrast, Subpopulations 2 and 3 are composed mainly of landraces that typically have higher genetic diversity, resulting from undergoing natural selection, which in general have rapid LD decay patterns

### 3.3 ASSOCIATION MAPPING FOR BLS RESISTANCE

The quantile-quantile (QQ) plot for MLM, MLMM, FarmCPU and BLINK methods conformed well with the expected distribution and did not show genomic inflation that could be brought about by the confounding effects of population structure (Figure 3). The deviation at the right tail end of the QQ plot of each method indicates evidence of association to BLS response. Using these four mixed linear methods for the GWAS, four significant marker-trait associations were consistently identified in at least two methods (Figure 4). These significant marker-trait associations include two associations mapped on chromosome 2H, and one each on chromosomes 5H, and 7H. The marker with the highest phenotypic variance explained (PVE) was BK_12, located at 25.87 Mbp on chromosome 2H, with a PVE of 21.00%. It is followed by 12_10306 at 365.80 Mbp on chromosome 5H, with 19.24% PVE (Table 2). Additional consistently significant markers include 11_10796 at 192.27 Mbp on chromosome 2H (3.70% PVE), and 12_30481 at 228.40 Mbp on chromosome 7H (8.37% PVE). Using the BLINK method, other significant MTAs that were detected include 12_30715 on chromosome 1H (4.59% PVE), 11_21125 on chromosome 2H (8.02% PVE), and SCRI_RS_237688 on chromosome 2H (5.66% PVE) (Table 2).

**Figure 3.**
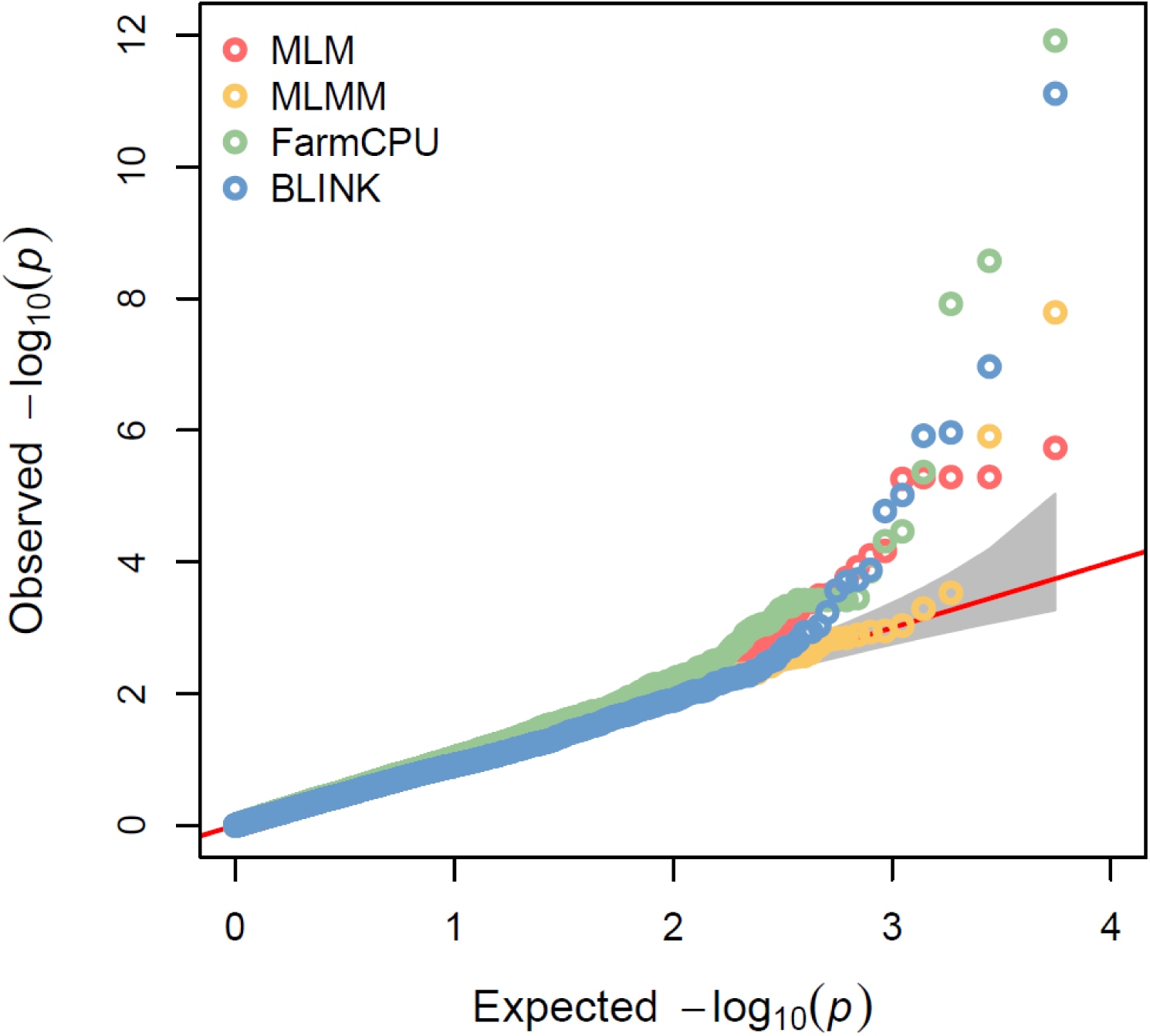
QQ plots of the four mixed models used in the GWAS analysis for BLS resistance on the barley mini-core panel including Mixed Linear Model (MLM), Multiple Loci Mixed Model (MLMM), Fixed and random model Circulating Probability Unification (FarmCPU), and Bayesian-information and Linkage-disequilibrium Iteratively Nested Keyway (BLINK).

**Figure 4.**
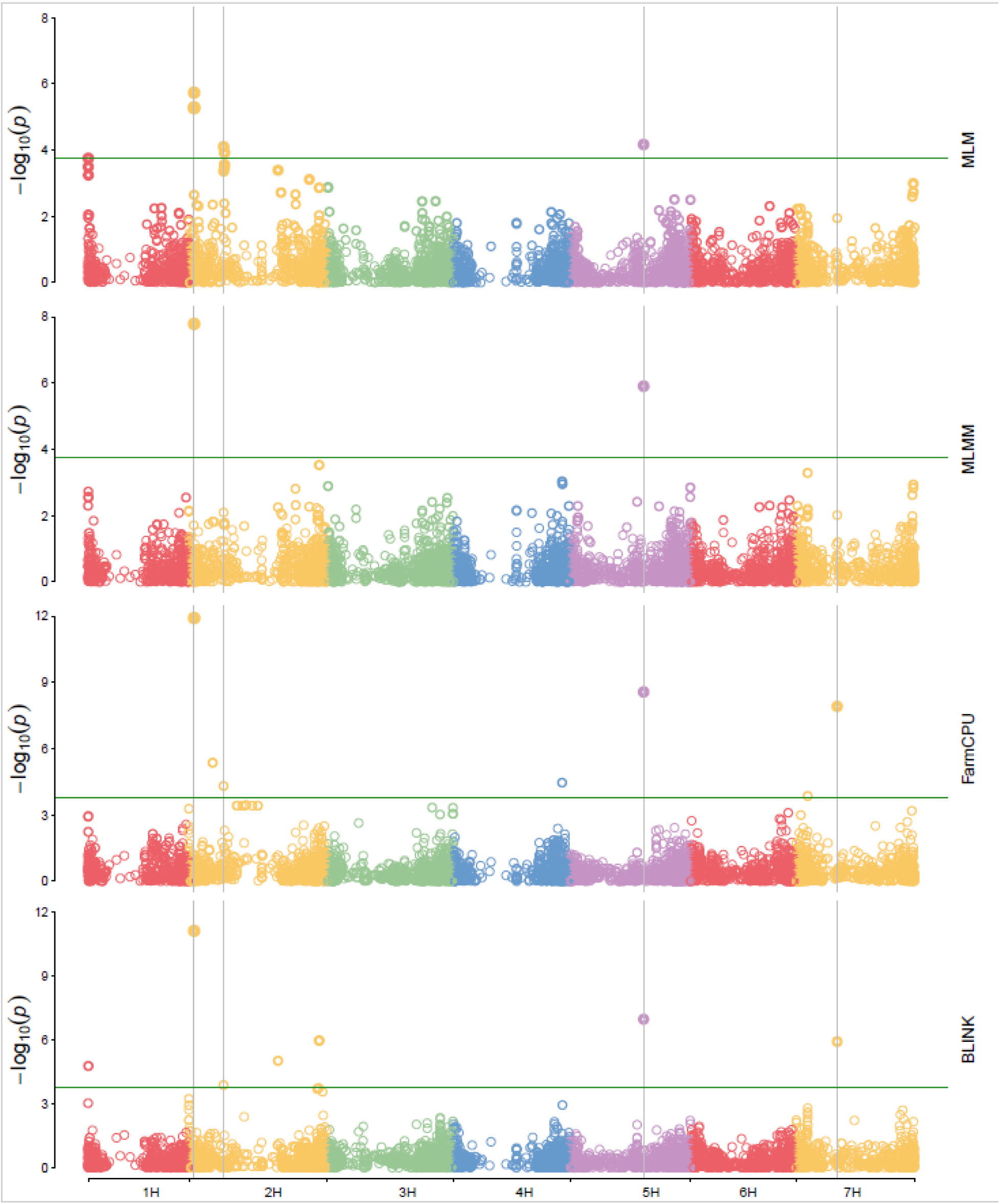
Manhattan plots of GWAS, from four different models, to identify markers that are potentially associated with BLS resistance. The solid lines indicate that significant markers were detected in at least two methods (including GLM model but not shown in photo).

**Table 2.**
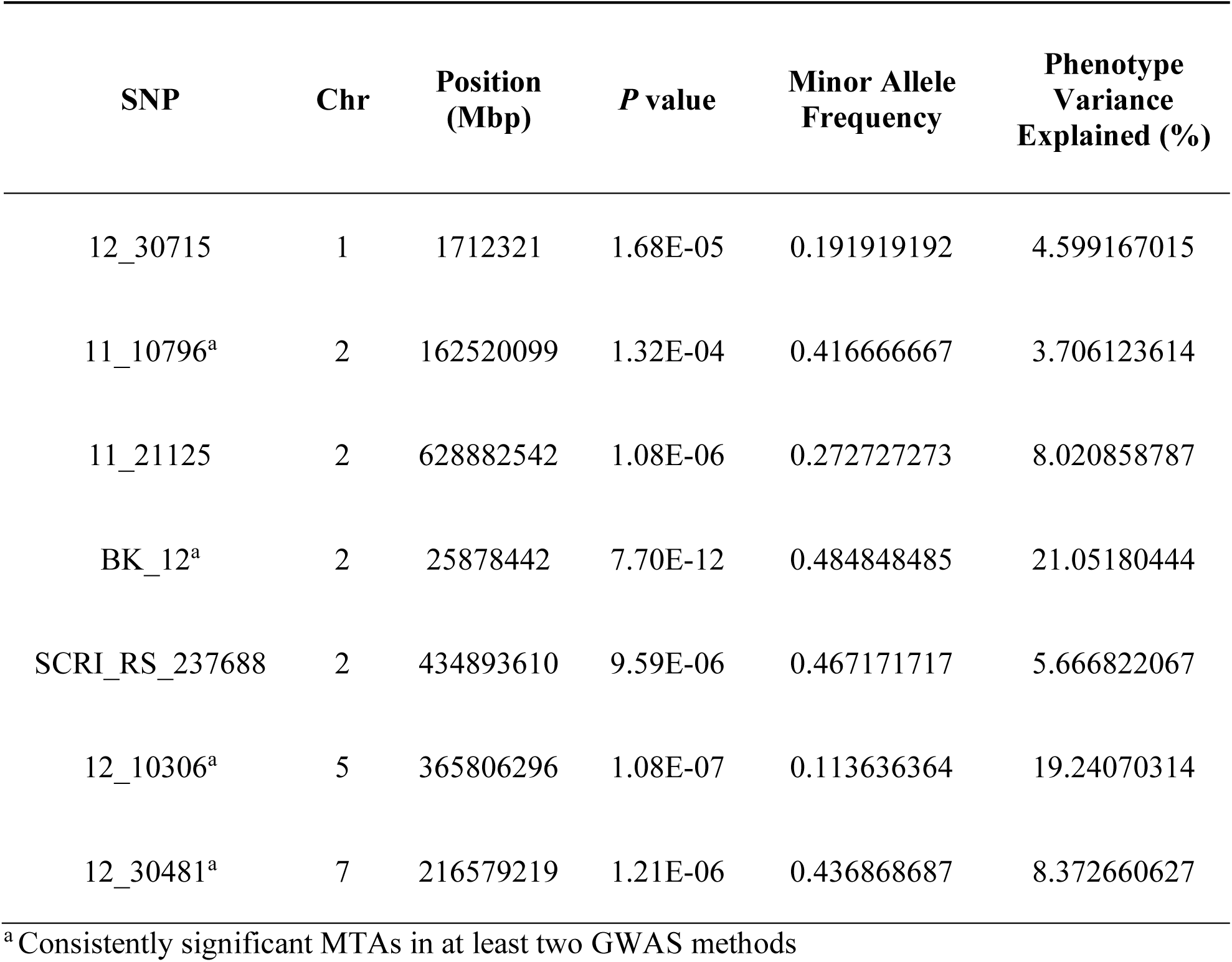
Significant MTAs in GWAS analysis (BLINK) for bacterial leaf streak resistance in the barley mini-core panel.

| SNP | Chr | Position (Mbp) | *P* value | Minor Allele Frequency | Phenotype Variance Explained (%) |
| --- | --- | --- | --- | --- | --- |
| 12_30715 | 1 | 1712321 | 1.68E-05 | 0.191919192 | 4.599167015 |
| 11_10796<sup>a</sup> | 2 | 162520099 | 1.32E-04 | 0.416666667 | 3.706123614 |
| 11_21125 | 2 | 628882542 | 1.08E-06 | 0.272727273 | 8.020858787 |
| BK_12<sup>a</sup> | 2 | 25878442 | 7.70E-12 | 0.484848485 | 21.05180444 |
| SCRI_RS_237688 | 2 | 434893610 | 9.59E-06 | 0.467171717 | 5.666822067 |
| 12_10306<sup>a</sup> | 5 | 365806296 | 1.08E-07 | 0.113636364 | 19.24070314 |
| 12_30481<sup>a</sup> | 7 | 216579219 | 1.21E-06 | 0.436868687 | 8.372660627 |

<sup>a</sup> Consistently significant MTAs in at least two GWAS methods

## 5 DISCUSSIONS

BLS disease of barley is a pernicious problem, and its economic implications may only worsen if no intervention is developed to manage it. With the absence of effective crop protection products against BLS in cereals, the use of resistant cultivars remains to be the most practical and economical method of control (Forster and Schaad 1988; Lux et al. 2023; Dusek et al. 2025). The susceptibility of current commercial varieties to BLS implies that current genetic pools in breeding programs lack sources of BLS resistance (Friskop et al. 2023). This compels us to explore diverse barley germplasms to identify potential donors of resistance. The investigation of Ritzinger et al. (2023a) on a large collection of breeding lines, landraces, and wild barley introgression lines revealed that the proportion of highly resistant lines was very low (4.3%). However, unlike the iCore set from the BCC, genetic redundancies were not evaluated on this set which may have amplified the proportion of susceptibility in the population. The BLS mini-core panel is a subset of the iCore collection which represents the global barley diversity. This panel provides an advantageous amount of genetic variation to maximize opportunities of identifying markers that are associated with QTLs responsible for BLS responses.

In the GWAS we conducted, five significant MTAs were consistently identified in at least two mixed model analyses. These associations are in chromosomes 2H, 5H, and 7H (Figure 4; Table 2). Most of the identified MTAs were also found in NLR-rich regions identified in the study of Li et al. (2021). Of the four MTAs that were consistently identified, BK_12 (25.87 Mbp) in chromosome 2H was the same marker found in the GWAS conducted by Ritzinger et al. (2023b) using different populations. Similar to their report, the marker BK_12, that coincides with the photoperiod response gene (*Ppd-H1*) of barley, also had the highest PVE across all MTAs identified to be associated with BLS response in this population. This observation fortifies the notion that maturity and BLS responses are related and that gene/s responsible for resistance could be tightly linked with the photoperiod response gene – explaining why most resistance is observed on lines with late maturity (Mizrak, 1985; Ritzinger et al. 2023a, b). In conjunction to this, the highly resistant accessions in this mini-core panel are predominantly two-rowed, and have consistently exhibited low disease infection levels over four years of observations (Table 1). In agreement with previous findings, six-rowed barley matures earlier than two-rowed barley and often shows more susceptibility to BLS. The majority of these resistant accessions are older breeding lines and cultivars, but several landraces also showed high resistance across four years of testing. This underscores the importance of exploring populations with further genetic diversities from current breeding pools to identify potential sources of BLS resistance. These accessions originated from diverse regions, including nine European countries, three African countries and three Asian countries, two from the Americas (Table 1). Due to their consistent resistant performance, these accessions are valuable genetic resources for advancing research on potential sources of BLS resistance gene/s.

In determining the candidate gene region in this GWAS, the critical threshold of r2 = 0.1 was used to identify the distance where LD of the mini-core panel has decayed. However, with subpopulations showing variability in LD decay patterns, it is important to remember that LD and its decay are specific to a population (Supplemental Figure S2) (Otyama, et al. 2019; Chao et al. 2010). Therefore, the respective distances at which LD decayed for each subpopulation must be considered as well in defining the candidate gene region for the respective subpopulations. LD of subpopulations 1 and 4 decayed at around 4.4 Mb, while subpopulations 2 and 3 decayed at approximately 0.4 Mb. The huge disparity between the distance at which LD decayed in these subpopulations may be affected by the differences in the accessions that comprise them.

In addition, the number of markers that were available for the population may have also limited the mapping resolution. It is important to note that if the associated markers of causal variants are not part of the panel, then the risk of missing the detection of important QTLs increases (Myles et al. 2009; Clark et al. 2005; Xu et al. 2017). The relatively limited number of markers available to use for mapping a structured population such as this mini-core panel might reduce the power to accurately identify potentially associated loci to BLS responses. Given the rapid LD decay in subpopulations 2 and 3, a larger set of markers may be needed to increase the mapping resolution and improve the ability to more accurately identify causal variants for BLS resistance. And while some MTAs from this GWAS align with previous studies, it may be prudent to re-genotype the panel using a more robust set of markers, such as the 50k iSelect SNP platform (Bayer et al. 2017), to enhance the resolution of the association mapping. Furthermore, recent advances in population diversity on *Xanthomonas translucens* pv. *translucens* revealed that the barley-specific BLS pathogen has three distinct clades with diverse effector sets (Curland et al. 2018; Heiden et al. 2023). These findings highlight the importance of looking into potential specific host-pathogen interactions where different sets of loci may be associated with different strains of the pathogen. For context, in this study one isolate from clade K1 was used to artificially inoculate, however, other clades could be present in the field.

In conclusion, through the GWAS conducted on the subset of the barley core collection, we identified markers that are significantly associated with BLS response, and we also verified the strong association of the *Ppd-H1* gene, through the marker BK_12 in chromosome 2H, to BLS resistance. The MTAs that we have reported in this study provide valuable insights and opportunities to explore potentially associated resistance loci. Additionally, the top-performing accessions are promising germplasms that may be evaluated to a greater extent for their BLS resistance across various environments or against different strains of the pathogen. Altogether, these findings enhance our understanding of host resistance to BLS, supporting global barley improvement efforts.

## ACKNOWLEDGMENTS

The work was supported by American Malting Barley Association. This research was supported by the American Malting Barley Association (AMBA) through the grant “Management and Innovative Research on Economically Important Barley Diseases”. Additional support was provided by the USDA–ARS Barley Pest Research Initiative under the project “Integrated Management of Barley Foliar Pathogens Including Spot Blotch, Bacterial Leaf Streak, and Net Blotch”, and by the North Dakota State Board of Agricultural Research and Education (SBARE) through the project “Genetic Diversity and Virulence of Barley Bacterial Leaf Streak Pathogen Populations”.

## SUPPLEMENTAL MATERIAL

**Supplemental Figure S1.** (A) Population structure analysis showed peaks at Δk=2 and Δk=4, where the former effectively divides the population based on the row type while the latter clustered the accessions based on their respective regions of origin; (B) Ancestry plot based on Δk=4 with a membership coefficient of >0.7.

**Supplemental Figure S2.** (A) LD of the mini-core panel shows rapid decay as distance increases. (B, C, D, E) LD of individual subpopulations exhibit variable decay patterns. Subpopulations 1 and 4 showed a steadier decay pattern compared to the rapid decay observed on LD of Subpopulations 2 and 3. (black lines show the LD decay pattern; blue broken lines show the critical threshold r2 = 0.1.

**Supplemental Figure S3.** Combined Manhattan plots of GWAS conducted on the mini-core panel (2013–2016) using the different mixed model methods, highlighting the significant MTAs identified across each chromosome.

**Supplemental Table S1.** List of accessions in the mini-core panel of the barley core collection, detailing their geographic origin, row type, improvement status, and disease severity scores.

**Supplemental Table S2.** Composition of each subpopulation based on improvement status.

**Supplemental Table S3.** MTAs identified through GWAS on the mini-core panel (2013-2016) using different mixed model methods.

## Abbreviations

BLINK: Bayesian-information and Linkage-disequilibrium Iteratively Nested Keyway
BLS: bacterial leaf streak
FarmCPU: Fixed and random Circulating Probability Unification
GWAS: genome-wide association study
LD: linkage disequilibrium
MLM: mixed linear model
MLMM: multi-locus mixed model
MTA: marker-trait association
PVE: phenotypic variance explained
QTL: quantitative trait loci
SNP: single nucleotide polymorphism
Xtt: *Xanthomonas translucens* pv. *translucens*

