## Supplementary Tables for "Genome-wide association study of host resistance to bacterial leaf streak in a subset of the world barley core collection"

**Supplementary Table S1** List of accessions in the mini-core panel of the barley core collection, detailing their geographic origin, row type, improvement status, and disease severity scores.

| Accession | Name | ORIGIN | Country | RECEIVED | Row Type | Improvement Status | BLS Score (BLUP) |
|---|---|---|---|---|---|---|---|
| PI 611526 | ELS6402-317 | Oromīya, Ethiopia | Ethiopia | 1964 | 6 | LANDRACE | 2.01 |
| PI 477775 | UNA8301 | Puno, Peru | Peru | 1983 | 6 | LANDRACE | 2.37 |
| PI 197103 | Shizuoka_Shirorokkaku1 | Sizuoka, Japan | Japan | 1951 | 6 | CULTIVAR | 2.41 |
| PI 87403 | SWANS_NECK | Lithuania | Lithuania | 1930 | 2 | UNCERTAIN | 2.56 |
| PI 184880 | Erectoides_No.23 | Skåne län, Sweden | Sweden | 1949 | 2 | BREEDING | 2.60 |
| PI 361695 | Lapinohra | Finland | Finland | 1971 | 6 | CULTIVAR | 2.61 |
| PI 190790 | Inumo_O | Korea, North | North Korea | 1950 | 6 | LANDRACE | 2.68 |
| CIho 6277 | Bulgarian347 | Bulgaria | Bulgaria | 1937 | 6 | UNCERTAIN | 2.76 |
| PI 306477 | 2734 | Romania | Romania | 1965 | 6 | UNCERTAIN | 2.89 |
| PI 243181 | MINERVA\|PI243181 | Gelderland, Netherlands | Netherlands | 1957 | 2 | CULTIVAR | 2.99 |
| PI 386407 | IAR/B/253 | Gash-Barka, Eritrea | Eritrea | 1974 | 2 | LANDRACE | 3.06 |
| PI 467460 | Parou | Greece | Greece | 1982 | 6 | CULTIVAR | 3.09 |
| PI 392491 | Welgevallen60-A-3-11 | Cape Province, South Africa | South Africa | 1974 | 2 | BREEDING | 3.12 |
| PI 306469 | 2724 | Romania | Romania | 1965 | 6 | UNCERTAIN | 3.13 |
| PI 392501 | Welgevallen65-31-36 | Cape Province, South Africa | South Africa | 1974 | 2 | BREEDING | 3.15 |
| PI 643247 | ND8763 | North Dakota, United States | United States | 2006 | 2 | BREEDING | 3.19 |
| PI 313113 | Europa | Baden-Wurttemberg, Germany | Germany | 1966 | 2 | CULTIVAR | 3.25 |
| PI 328855 | Mansholts_Tweerijige | Groningen, Netherlands | Netherlands | 1968 | 2 | CULTIVAR | 3.28 |
| PI 386759 | IAR/B/400 | Ethiopia | Ethiopia | 1974 | 6 | LANDRACE | 3.32 |
| PI 467808 | PERFEKTA\|PI467808 | Austria | Austria | 1982 | 2 | CULTIVAR | 3.32 |
| PI 467419 | Pisarecky | Czechoslovakia | Czechoslovakia | 1982 | 2 | CULTIVAR | 3.34 |
| PI 265465 | Paavo | Finland | Finland | 1960 | 2 | CULTIVAR | 3.40 |
| PI 467438 | SANDOMIERSKI\|PI467438 | Poland | Poland | 1982 | 2 | CULTIVAR | 3.41 |
| CIho 14280 | CI14280 | Táchira, Venezuela | Venezuela | 1925 | 2 | UNCERTAIN | 3.46 |
| PI 361676 | 6Radet_Nogen | Denmark | Iraq | 1971 | 6 | UNCERTAIN | 3.49 |
| PI 321844 | Kupljenik7 | Slovenia | Slovenia | 1967 | 2 | CULTIVAR | 3.58 |
| CIho 11789 | Djeddah | Saudi Arabia | Saudi Arabia | 1960 | 2 | UNCERTAIN | 3.63 |
| PI 356711 | MOR11/6 | Morocco | Morocco | 1970 | 6 | LANDRACE | 3.67 |
| PI 69421 | 6995 | Hebei Sheng, China | China | 1926 | 6 | LANDRACE | 3.70 |
| PI 370782 | 142A | Graubünden, Switzerland | Switzerland | 1971 | 2 | LANDRACE | 3.71 |
| PI 467354 | Finnebygg | Norway | Norway | 1982 | 6 | CULTIVAR | 3.71 |
| PI 371100 | 1065B | Valais, Switzerland | Switzerland | 1971 | 2 | LANDRACE | 3.74 |
| PI 344903 | PI344903 | Serbia | Serbia | 1969 | 2 | LANDRACE | 3.76 |
| PI 467815 | Etu | Finland | Finland | 1982 | 6 | CULTIVAR | 3.80 |
| CIho 2236 | REIDS_TRIUMPH | Idaho, United States | United States | 1920 | 6 | UNCERTAIN | 3.81 |
| PI 434760 | QB704.3 | Québec, Canada | Canada | 1979 | 6 | BREEDING | 3.84 |
| PI 290363 | Schonfinower | Pest, Hungary | Hungary | 1963 | 6 | UNCERTAIN | 3.86 |
| PI 611562 | Obskoi | Novosibirskaja oblast', Russian Federation | Russia | 1999 | 2 | CULTIVAR | 3.90 |
| PI 573591 | Podolskij8 | Vinnytska oblast, Ukraine | Ukraine | 1992 | 2 | CULTIVAR | 3.90 |
| PI 343725 | Crusader | Groningen, Netherlands | Netherlands | 1969 | 2 | CULTIVAR | 3.90 |
| PI 382982 | GAW87-1 | Tigray, Ethiopia | Ethiopia | 1973 | 6 | LANDRACE | 3.98 |
| PI 399491 | Diva | Zuid-Holland, Netherlands | Netherlands | 1975 | 2 | CULTIVAR | 3.99 |
| PI 428394 | Arabic_Akhdar | Syria | Syria | 1978 | 2 | UNCERTAIN | 4.01 |
| | Belorusskij18 | Horad Minsk, Belarus | Belarus | 1972 | 2 | CULTIVAR | 4.03 |
| | Varunda | Gelderland, Netherlands | Netherlands | 1976 | 2 | CULTIVAR | 4.09 |
| PI 525199 | Beatrice | Île-de-France, France | France | 1988 | 2 | CULTIVAR | 4.11 |
| | 2550 | Al Minūfīyah, Egypt | Egypt | 1988 | 2 | UNCERTAIN | 4.12 |

| | | | | | | | |
|---|---|---|---|---|---|---|---|
| | Elliot | United Kingdom | United Kingdom | 1993 | 2 | CULTIVAR | 4.19 |
| | Nolc_Dregeruv_Bohemia | Czechoslovakia | Czech Republic | 1968 | 2 | CULTIVAR | 4.19 |
| | Novoomskii | Omskaja oblast', Russian Federation | Russia | 1999 | 2 | CULTIVAR | 4.20 |
| | Black_Roummani | Al Ḩasakah, Syria | Syria | 1949 | 2 | LANDRACE | 4.21 |
| | Akme_Futter | Slovenia | Slovenia | 1967 | 2 | CULTIVAR | 4.22 |
| | Rupe | South Island, New Zealand | New Zealand | 1985 | 2 | CULTIVAR | 4.25 |
| | HOR4181 | Turkmenistan | Turkmenistan | 1968 | 6 | LANDRACE | 4.27 |
| | NUDIDEFICIENS | Russian Federation | Russia | 1924 | 2 | LANDRACE | 4.27 |
| | Gloire_du_Velay | Île-de-France, France | France | 1982 | 2 | CULTIVAR | 4.28 |
| | PI436143 | La Araucanía, Chile | Chile | 1979 | 2 | LANDRACE | 4.28 |
| | 21d | Īlām, Iran | Iran | 1963 | 6 | LANDRACE | 4.29 |
| | Primus_II | Skåne län, Sweden | Sweden | 1960 | 2 | CULTIVAR | 4.32 |
| | Pulawski | Krakow, Poland | Poland | 1938 | 2 | CULTIVAR | 4.32 |
| | HATVANI308\|PI328835 | Heves, Hungary | Hungary | 1968 | 2 | CULTIVAR | 4.32 |
| | PI447207 | Aragón, Spain | Spain | 1980 | 2 | UNCERTAIN | 4.32 |
| | FL7138A3-102 | Florida, United States | United States | 1978 | 6 | BREEDING | 4.33 |
| | Antarctica01 | Brazil | Brazil | 1982 | 2 | CULTIVAR | 4.36 |
| | Israeli121 | Israel | Israel | 1964 | 2 | CULTIVAR | 4.37 |
| | Weibull5492 | Skåne län, Sweden | Sweden | 1963 | 2 | BREEDING | 4.38 |
| | SN-33 | Samegrelo-Zemo Svaneti, Georgia | Georgia | 1991 | 2 | LANDRACE | 4.39 |
| | REX | Denmark | Denmark | 1920 | 2 | CULTIVAR | 4.41 |
| | Triumf | South Moravia, Czech Republic | Czech Republic | 1960 | 2 | CULTIVAR | 4.42 |
| | Kosmos | Poland | Poland | 1982 | 2 | CULTIVAR | 4.45 |
| | 4008 | Mongolia | Mongolia | 1924 | 6 | LANDRACE | 4.46 |
| | Opavsky_Kneifel | Czechoslovakia | Czechoslovakia | 1982 | 2 | CULTIVAR | 4.47 |
| | Nitransky_Exportni | West Slovakia, Slovakia | Slovakia | 1978 | 2 | CULTIVAR | 4.52 |
| | Long_Glumes | Denmark | Denmark | 1971 | 2 | UNCERTAIN | 4.52 |
| | Welgevallen65-41-7 | Cape Province, South Africa | South Africa | 1974 | 2 | BREEDING | 4.55 |
| | Bikai | Al Biqā‘, Lebanon | Lebanon | 1949 | 2 | LANDRACE | 4.56 |
| | HOR1007 | Thessalía, Greece | Greece | 1962 | 6 | UNCERTAIN | 4.56 |
| | RPB421 | Lincolnshire, United Kingdom | United Kingdom | 1984 | 2 | BREEDING | 4.56 |
| | Hodoninsky_Kvas | East Slovakia, Slovakia | Slovakia | 1948 | 2 | CULTIVAR | 4.58 |
| | Pukako | South Island, New Zealand | New Zealand | 1985 | 2 | CULTIVAR | 4.58 |
| | Hankkija673 | Finland | Finland | 1975 | 6 | CULTIVAR | 4.59 |
| | Ilineskij43 | Vinnytska oblast, Ukraine | Ukraine | 1965 | 2 | CULTIVAR | 4.59 |
| | PI344944 | Pljevlja, Montenegro | Montenegro | 1969 | 2 | LANDRACE | 4.61 |
| | Welgevallen60-3-88A | Cape Province, South Africa | South Africa | 1974 | 2 | BREEDING | 4.61 |
| | QB139.8 | Québec, Canada | Canada | 1975 | 6 | BREEDING | 4.61 |
| | HOR1009 | Pelopónnisos, Greece | Greece | 1968 | 2 | LANDRACE | 4.64 |
| | 10250 | Turkistan | Turkistan | 1926 | 2 | BREEDING | 4.65 |
| | HOR4097 | Turkmenistan | Turkmenistan | 1968 | 6 | LANDRACE | 4.65 |
| | 8389 | Jilin Sheng, China | China | 1926 | 6 | LANDRACE | 4.65 |
| | Motan | Occitanie, France | France | 1993 | 6 | CULTIVAR | 4.66 |
| | TX01D196 | Texas, United States | United States | 2003 | 6 | BREEDING | 4.70 |
| | TX99D923 | Texas, United States | United States | 2003 | 6 | BREEDING | 4.70 |
| | Welgevallen65-55-3 | Cape Province, South Africa | South Africa | 1974 | 2 | BREEDING | 4.70 |
| | Harpoon | Lincolnshire, United Kingdom | United Kingdom | 1984 | 2 | CULTIVAR | 4.71 |
| | BUCHER | Diyālá, Iraq | Ukraine | 1920 | 6 | LANDRACE | 4.71 |
| | 4041 | Mongolia | Mongolia | 1924 | 6 | LANDRACE | 4.72 |

| | | | | | | | |
|---|---|---|---|---|---|---|---|
| | PI94834 | Kharkivska oblast, Ukraine | Ukraine | 1930 | 2 | LANDRACE | 4.74 |
| | Maroccaine0187 | Morocco | Morocco | 1949 | 6 | CULTIVAR | 4.75 |
| | QB714.5 | Québec, Canada | Canada | 1979 | 6 | BREEDING | 4.76 |
| | MOR12/5 | Morocco | Morocco | 1970 | 6 | LANDRACE | 4.79 |
| | PI270618 | Puno, Peru | Peru | 1961 | 6 | LANDRACE | 4.79 |
| | SN-11 | Samegrelo-Zemo Svaneti, Georgia | Georgia | 1991 | 2 | LANDRACE | 4.85 |
| | Chae-Rae-Baec | Korea, South | South Korea | 1950 | 6 | LANDRACE | 4.87 |
| | Fundulea245/64 | Călărași, Romania | Romania | 1969 | 6 | CULTIVAR | 4.88 |
| | MCU3674 | Cundinamarca, Colombia | Colombia | 1974 | 6 | UNCERTAIN | 4.93 |
| | 3998 | Mongolia | Mongolia | 1924 | 6 | LANDRACE | 4.94 |
| | KHARTUM | Khartoum, Sudan | Sudan | 1917 | 6 | UNCERTAIN | 4.98 |
| | CI14291 | Sichuan Sheng, China | China | 1925 | 6 | LANDRACE | 5.01 |
| | Baku | Iran | Iran | 1910 | 2 | LANDRACE | 5.01 |
| | Jao_Charpalu | Sar-e Pul, Afghanistan | Afghanistan | 1940 | 6 | LANDRACE | 5.02 |
| | ND4998 | North Dakota, United States | United States | 2006 | 2 | BREEDING | 5.04 |
| | 69Z108.179 | Salzburg, Austria | Austria | 1970 | 2 | LANDRACE | 5.07 |
| | JM-3988 | West Bank, Israel | Israel | 1991 | 6 | LANDRACE | 5.11 |
| | STB76 | Lincolnshire, United Kingdom | United Kingdom | 1981 | 6 | BREEDING | 5.14 |
| | PI94806 | Russian Federation | Russia | 1930 | 2 | LANDRACE | 5.15 |
| | DJEBALI\|CIHO15291 | La Manouba, Tunisia | Tunisia | 1972 | 6 | LANDRACE | 5.16 |
| | TX01D292 | Texas, United States | United States | 2003 | 6 | BREEDING | 5.16 |
| | TX01D397 | Texas, United States | United States | 2003 | 6 | BREEDING | 5.18 |
| | Sandra | Île-de-France, France | France | 1978 | 2 | CULTIVAR | 5.20 |
| | C93 | North-West Frontier, Pakistan | Pakistan | 1976 | 6 | LANDRACE | 5.28 |
| | Martonvasari_MK-175 | Budapest, Hungary | Hungary | 1990 | 2 | CULTIVAR | 5.29 |
| | MENUET\|PI467789 | Netherlands | Netherlands | 1982 | 2 | CULTIVAR | 5.30 |
| | Hinowi | Egypt | Egypt | 1923 | 6 | UNCERTAIN | 5.31 |
| | Cerise | Lincolnshire, United Kingdom | United Kingdom | 1984 | 2 | CULTIVAR | 5.31 |
| | Baitori_No.1 | Kyôto, Japan | Japan | 1949 | 6 | CULTIVAR | 5.32 |
| | MCU3485 | Cundinamarca, Colombia | Colombia | 1974 | 6 | UNCERTAIN | 5.32 |
| | 3125 | Lara, Venezuela | Venezuela | 1935 | 6 | LANDRACE | 5.36 |
| | 5195 | Hormozgān, Iran | Iran | 1940 | 6 | LANDRACE | 5.36 |
| | ORZO_COLLESANO1921 | Puglia, Italy | Italy | 1922 | 6 | BREEDING | 5.36 |
| | ND16680 | North Dakota, United States | United States | 2006 | 2 | BREEDING | 5.39 |
| | B32A | Purwanchal, Nepal | Nepal | 1978 | 6 | LANDRACE | 5.40 |
| | Vantmore | Manitoba, Canada | Canada | 1954 | 6 | CULTIVAR | 5.47 |
| | Taiju_Omugi | Hokkaidô, Japan | Japan | 1950 | 6 | LANDRACE | 5.49 |
| | ATTIKI\|PI467797 | Macedonia, Greece | Greece | 1982 | 6 | CULTIVAR | 5.49 |
| | II-20 | Croatia | Croatia | 1960 | 6 | UNCERTAIN | 5.52 |
| | CI13353 | Āmara, Ethiopia | Ethiopia | 1964 | 2 | LANDRACE | 5.54 |
| | JAO\|PI176144 | Uttarakhand, India | India | 1949 | 6 | LANDRACE | 5.55 |
| | ISOGENIC:2-ROW_LARGE_LATERAL_LATE | Montana, United States | United States | 1980 | 2 | GENETIC | 5.55 |
| | CAVEDA | Azores, Portugal | Portugal | 1904 | 6 | UNCERTAIN | 5.59 |
| | CI7503 | Puebla, Mexico | Mexico | 1947 | 6 | LANDRACE | 5.59 |
| | II-831-4B-1B-1-5B | Cundinamarca, Colombia | Colombia | 1957 | 6 | BREEDING | 5.62 |
| | S-42 | Āmara, Ethiopia | Ethiopia | 1962 | 6 | LANDRACE | 5.65 |
| | SN-84a | Samegrelo-Zemo Svaneti, Georgia | Georgia | 1991 | 2 | LANDRACE | 5.65 |
| | Surprise-II-750-3B-1B-1-7T | Cundinamarca, Colombia | Colombia | 1957 | 6 | BREEDING | 5.67 |
| | Ali | Shaanxi Sheng, China | China | 1992 | 6 | CULTIVAR | 5.78 |

| | | | | | | | |
|---|---|---|---|---|---|---|---|
| | PI264907 | Nótio Aigaío, Greece | Greece | 1960 | 6 | LANDRACE | 5.79 |
| | BELADI\|PI283433 | Cyprus | Cyprus | 1962 | 6 | LANDRACE | 5.86 |
| | PI270641 | Puno, Peru | Peru | 1961 | 6 | LANDRACE | 5.87 |
| | C.A.N.1105 | Ontario, Canada | Canada | 1935 | 6 | BREEDING | 5.94 |
| | CI5329 | Idaho, United States | United States | 1929 | 6 | BREEDING | 5.99 |
| | ND10419 | North Dakota, United States | United States | 2006 | 2 | BREEDING | 6.00 |
| | PERUVIAN | Cusco, Peru | Peru | 1917 | 6 | UNCERTAIN | 6.01 |
| | Kaljan | Afghanistan | Afghanistan | 1937 | 6 | LANDRACE | 6.01 |
| | NIGRUM | Iraq | Iraq | 1921 | 6 | LANDRACE | 6.08 |
| | Afghanistan600 | Brașov, Romania | Romania | 1971 | 6 | UNCERTAIN | 6.14 |
| | CARRE42 | Algeria | Algeria | 1923 | 6 | UNCERTAIN | 6.17 |
| | Sudan | Lisboa, Portugal | Portugal | 1938 | 6 | UNCERTAIN | 6.18 |
| | 10246 | Turkistan | Turkistan | 1926 | 6 | BREEDING | 6.23 |
| | CI7492 | Hidalgo, Mexico | Mexico | 1947 | 6 | LANDRACE | 6.34 |
| | II-11986-2b-3t-3b | Cundinamarca, Colombia | Colombia | 1968 | 6 | BREEDING | 6.40 |
| | 3839 | İzmir, Turkey | Turkey | 1948 | 2 | LANDRACE | 6.43 |
| | TX99D684 | Texas, United States | United States | 2003 | 6 | BREEDING | 6.43 |
| | NB5 | Israel | Israel | 1964 | 6 | CULTIVAR | 6.44 |
| | RNB-373 | Purwanchal, Nepal | Nepal | 1992 | 6 | LANDRACE | 6.46 |
| | HOR4101 | Uzbekistan | Uzbekistan | 1968 | 6 | LANDRACE | 6.47 |
| | CI14286 | Chile | Chile | 1924 | 6 | LANDRACE | 6.49 |
| | Barquis | Lara, Venezuela | Venezuela | 1914 | 6 | LANDRACE | 6.53 |
| | Urbush | Balochistan, Pakistan | Pakistan | 1992 | 6 | LANDRACE | 6.53 |
| | Heine_FI0367/53 | Niedersachsen, Germany | Germany | 1955 | 6 | BREEDING | 6.58 |
| | Nabawi | Jordan | Jordan | 1955 | 6 | UNCERTAIN | 6.64 |
| | MCU3523 | Cundinamarca, Colombia | Colombia | 1974 | 6 | UNCERTAIN | 6.68 |
| | DJEBALI\|CIHO15355 | Tunisia | Turkistan | 1972 | 6 | LANDRACE | 6.76 |
| | CI6496 | Xizang Zizhiqu, China | China | 1938 | 6 | UNCERTAIN | 6.79 |
| | Mutant_ERT7A | Buenos Aires, Argentina | Argentina | 1969 | 2 | BREEDING | 6.80 |
| | 4064 | Mongolia | Mongolia | 1924 | 6 | LANDRACE | 6.82 |
| | Moiris | Mary, Turkmenistan | Turkmenistan | 1919 | 6 | UNCERTAIN | 6.82 |
| | PAPHITICO\|PI283434 | Cyprus | Cyprus | 1962 | 6 | CULTIVAR | 6.88 |
| | TX01D148 | Texas, United States | United States | 2003 | 6 | BREEDING | 6.88 |
| | DJEBALI\|CIHO15290 | La Manouba, Tunisia | Tunisia | 1972 | 6 | LANDRACE | 6.89 |
| | PI343713 | Ar Riyāḑ, Saudi Arabia | Saudi Arabia | 1969 | 6 | UNCERTAIN | 6.92 |
| | PI447051 | Aragón, Spain | Spain | 1980 | 2 | BREEDING | 6.95 |
| | CAPE\|PI49154 | Matabeleland North, Zimbabwe | Zimbabwe | 1920 | 6 | UNCERTAIN | 6.97 |
| | Orge071 | Morocco | Morocco | 1962 | 6 | CULTIVAR | 7.05 |
| | IV/156 | North Macedonia | Macedonia | 1977 | 6 | LANDRACE | 7.08 |
| | PI84314 | Xorazm, Uzbekistan | Uzbekistan | 1930 | 6 | LANDRACE | 7.15 |
| | 6089 | Kābul, Afghanistan | Afghanistan | 1925 | 6 | LANDRACE | 7.18 |
| | TX02D285 | Texas, United States | United States | 2003 | 6 | BREEDING | 7.41 |
| | 1271 | Egypt | Egypt | 1988 | 6 | UNCERTAIN | 7.44 |
| | PI268255 | Fārs, Iran | Iran | 1960 | 2 | LANDRACE | 7.44 |
| | RNB-29 | Madhyamanchal, Nepal | Nepal | 1992 | 6 | LANDRACE | 7.50 |
| | 2606 | Kütahya, Turkey | Turkey | 1969 | 2 | LANDRACE | 7.50 |
| | Dneprovskii425 | Dnipropetrovska oblast, Ukraine | Slovakia | 1999 | 2 | CULTIVAR | 7.50 |
| | Welgevallen62-2-14-1 | Cape Province, South Africa | South Africa | 1974 | 2 | BREEDING | 7.60 |
| | Heran | Czech Republic | Czech Republic | 1997 | 2 | CULTIVAR | 7.66 |

| | | | | | | | |
|---|---|---|---|---|---|---|---|
| | Magnif132 | Buenos Aires, Argentina | Argentina | 1968 | 2 | CULTIVAR | 7.70 |
| PI 250775 | K710 | Kandahār, Afghanistan | Afghanistan | 1958 | 6 | LANDRACE | 7.76 |
| | Kaputar | New South Wales, Australia | Australia | 1994 | 2 | CULTIVAR | 7.77 |
| PI 320219 | C.P.I.28369 | Western Australia, Australia | Australia | 1967 | 6 | UNCERTAIN | 7.96 |
