## Supplementary Figures for "Genome-wide association study of host resistance to bacterial leaf streak in a subset of the world barley core collection"

A.

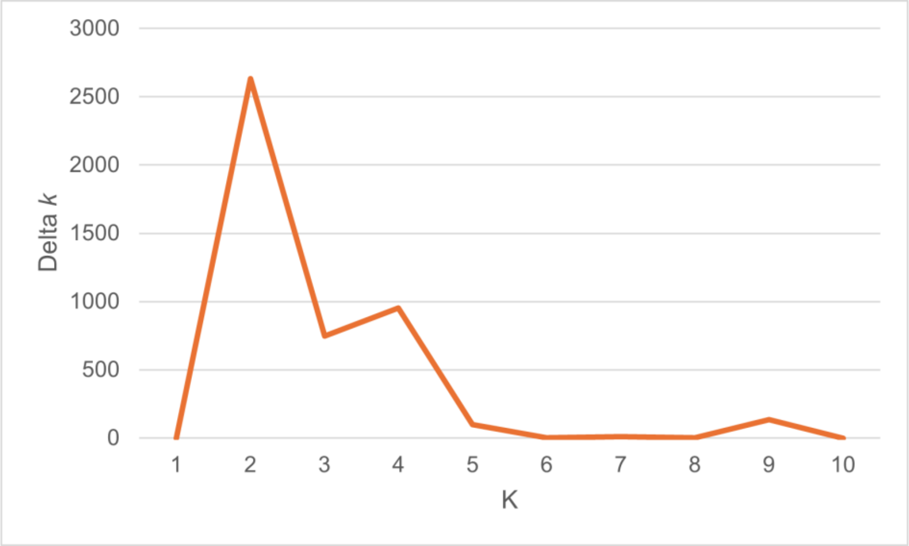


B.

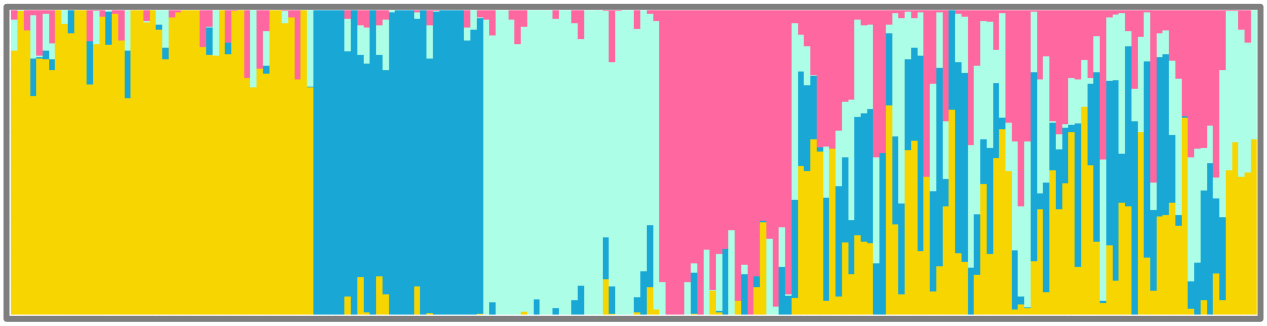

Supplementary Figure S1. (A) Population structure analysis showed peaks at Δ*k*=*2* and Δ*k*=*4*, where the former effectively divides the population based on the row type while the latter clustered the accessions based on their respective regions of origin; (B) Ancestry plot based on Δ*k*=4 with a membership coefficient of >0.7.

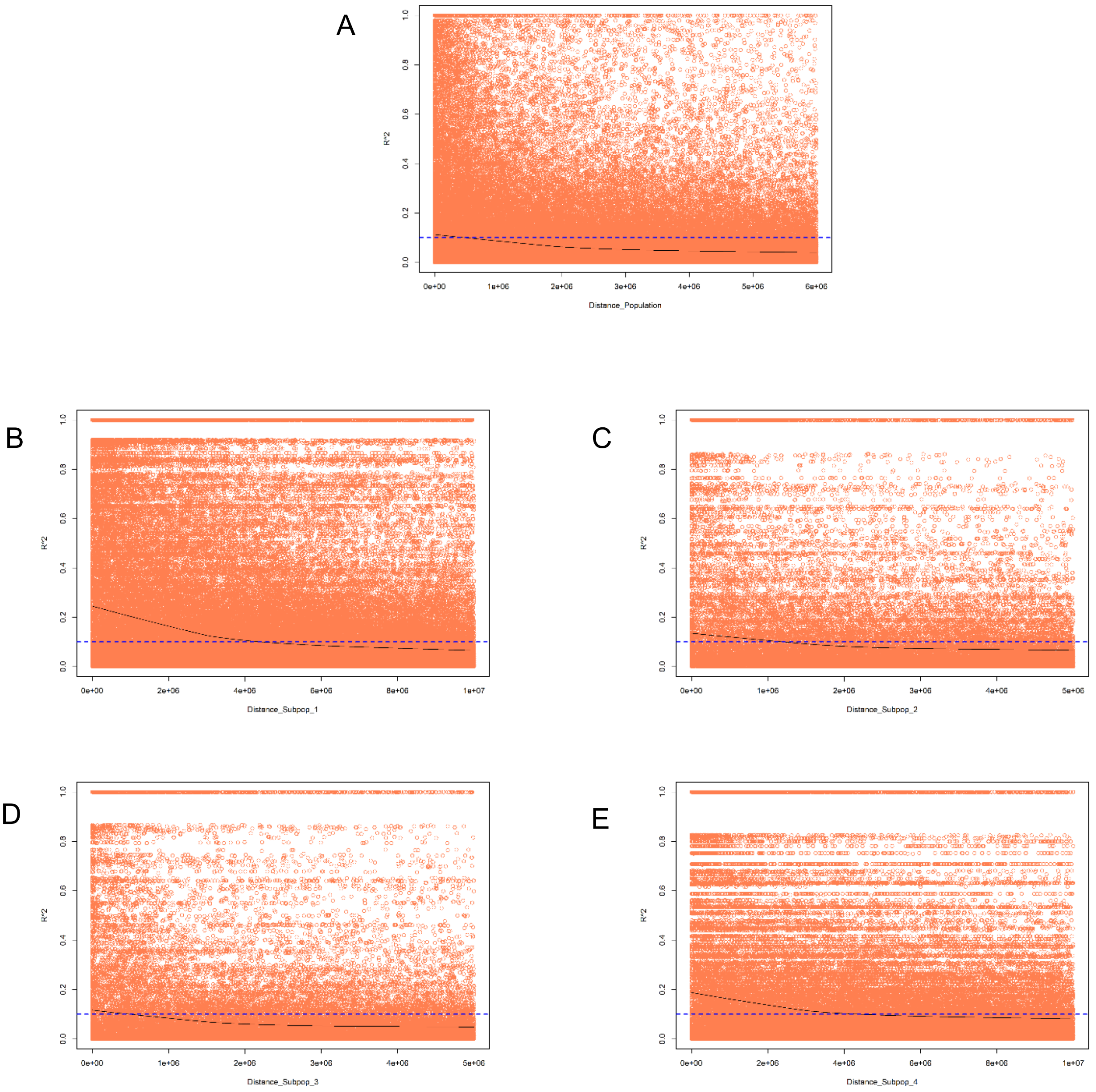


Supplementary Figure S2. (A) LD of the mini-core panel shows rapid decay as distance increases. (B, C, D, E) LD of individual subpopulations exhibit variable decay patterns. Subpopulations 1 and 4 showed a steadier decay pattern compared to the rapid decay observed on LD of Subpopulations 2 and 3. (black lines show the LD decay pattern; blue broken lines show the critical threshold $r^2 = 0.1$.

Supplementary Figure S3. Combined Manhattan plots of GWAS conducted on the mini-core panel (2013–2016) using the different mixed model methods, highlighting the significant MTAs identified across each chromosome.

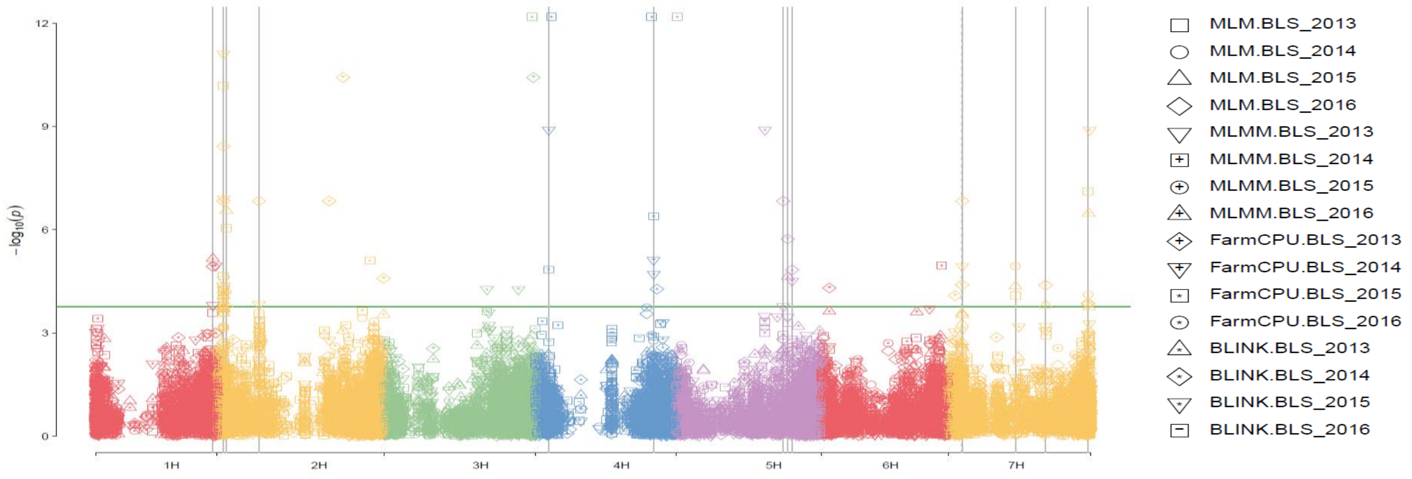
